# Investigation Gene body methylation is under selection in *Arabidopsis thaliana*

**DOI:** 10.1101/2020.09.04.283333

**Authors:** Aline Muyle, Jeffrey Ross-Ibarra, Danelle K. Seymour, Brandon S. Gaut

## Abstract

In plants, mammals and insects, some genes are methylated in the CG dinucleotide context, a phenomenon called gene body methylation. It has been controversial whether this phenomenon has any functional role. Here, we took advantage of the availability of 876 leaf methylomes in *Arabidopsis thaliana* to characterize the population frequency of methylation at the gene level and estimated the site-frequency spectrum of allelic states (epialleles). Using a population genetics model specifically designed for epigenetic data, we found that genes with ancestral gene body methylation are under significant selection to remain methylated. Conversely, all genes taken together were inferred to be under selection to be unmethylated. The estimated selection coefficients were small, similar to the magnitude of selection acting on codon usage. We also estimated that *A. thaliana* is losing gene body methylation three-fold more rapidly than gaining it, which could be due to a recent reduction in the efficacy of selection after a switch to selfing. Finally, we investigated the potential function of gene body methylation through its link with gene expression level. Across genes with polymorphic methylation states, the expression of gene body methylated alleles was consistently and significantly higher than unmethylated alleles. Although it is difficult to disentangle genetic from epigenetic effects, our work suggests that gbM has a small but measurable effect on fitness, perhaps due to its association to a phenotype like gene expression.

## Introduction

Cytosine DNA methylation is a type of epigenetic mark in which a methyl group is added to the 5^th^ carbon of cytosines. In plants, it can occur in three sequence contexts - CG, CHG and CHH (where H stands for A, T or C) – but levels and patterns of DNA methylation vary among genomic regions. In flowering plants, repetitive regions tend to be methylated in all three contexts, where methylation has a well-established repressive function on transposable elements (TEs) and regulatory elements (Luo *et al*. 2018; Schmitz *et al*. 2019). In contrast, exons in plants, insects and mammals are sometimes methylated only in the CG context. This gene body methylation (gbM) can be found within moderately and constitutively expressed housekeeping genes (Zhang *et al*. 2006; Neri *et al*. 2017; Schmitz *et al*. 2019) and is linked to active transcription in plants (Zhang *et al*. 2006; Zilberman *et al*. 2007; Cokus *et al*. 2008; Lister *et al*. 2008). However, it is not yet clear if gbM has a function, because the study of mutants deprived of gbM has failed to reveal a clear effect on phenotype (Teixeira and Colot 2009; Bewick and Schmitz 2017; Zilberman 2017).

The mechanisms responsible for the establishment of gbM in plants have recently been clarified, due in large part to studies in *Eutrema salsugineum*, a close relative of *Arabidopsis thaliana* that lacks both gbM and the *CHROMOMETHYLASE 3* (*CMT3*) gene (Bewick *et al*. 2016). The CMT3 protein has previously been shown to be involved in a self-reinforcing feedback loop: the histone mark H3K9me2 is recognized by CMT3 which then *de novo* methylates nearby cytosines in the CHG context and in turn leads to H3K9 methylation (Kawashima and Berger 2014). The deposition of CHG methylation typically suppresses transcription, but it is removed within transcribed genic regions by *INCREASED IN BONSAI METHYLATION 1 (IBM1)* (Saze *et al*. 2008; Miura *et al*. 2009).

The critical role of CMT3 in gbM establishment is supported by the facts that not just one but two Brassicaceae species have independently lost *CMT3* and both lack gbM (Bewick *et al*. 2016; Niederhuth *et al*. 2016). Moreover, transgenic reinsertion of CMT3 into *E. salsugineum* re-establishes genic methylation in all three contexts in a subset of genes that tend to be orthologous to gbM genes in *A. thaliana* (Wendte *et al*. 2019). This subset of genes has been called ‘CHG-gain’ genes (Wendte *et al*. 2019), and remarkably, these genes remained methylated only in the CG context following the loss of the *CMT3* transgene (Wendte *et al*. 2019). It remains unclear how CMT3 (and/or H3K9me2) is directed to a specific subset of genes for *de novo* DNA methylation and how these CHG-gain genes also become *de novo* methylated in the CG and CHH contexts (Wendte *et al*. 2019), but *cmt3* mutants in *A. thaliana* clearly demonstrate that CMT3 does not affect the maintenance of gbM once it is established (Stroud *et al*. 2013). Once CG methylation is established, it is maintained by METHYLTRANSFERASE 1 (MET1), which adds a methyl group on the symmetrical CG dinucleotide of a complementary DNA strand during cell duplication (Kawashima and Berger 2014). Maintenance by MET1 is an inherently error-prone process, as illustrated by epimutation accumulation in *A. thaliana* (Becker *et al*. 2011; Schmitz *et al*. 2011; van der Graaf *et al*. 2015). The accumulation of these epimutations over time illustrates that CG methylation is heritable.

Although gbM is widespread across species and relatively common within a genome - it is found for example in ∼20% of *A. thalia*na genes (Takuno and Gaut 2012) - it remains unclear whether gbM is functionally relevant (Teixeira and Colot 2009; Bewick and Schmitz 2017; Zilberman 2017). The question of its potential function has focused on three interrelated hypotheses. The first is that gbM affects gene expression. This hypothesis is supported by the fact that gbM genes exhibit a positive correlation between methylation and expression levels across genes (Zhang *et al*. 2006; Zilberman *et al*. 2007; Takuno and Gaut 2012), suggesting either that gbM might cause higher expression or, conversely, that active transcription drives gbM (Teixeira and Colot 2009). However, further tests of this association have led to contradictory results. For example, not all highly expressed genes have gbM in *A. thaliana* (Zhang *et al*. 2006; Zilberman *et al*. 2007), illustrating that any association is not absolute. The association has also been tested experimentally in epigenetic recombinant inbred lines (epiRILs) that were developed from the cross of a *met1* mutant and wild-type (WT) *A. thaliana*, followed by eight generations of inbreeding (Reinders *et al*. 2009). The resulting epiRILs had a mosaic methylome, with regions that have normal CG methylation derived from the WT parent and other regions derived from the *met1* mutant that originally lacked gbM. Analysis of gene expression in these lines detected no significant changes in the *met1* derived regions of epiRILs compared to orthologous WT regions (Bewick *et al*. 2016). Moreover, the epiRILs did not reestablish the original pattern of gbM after eight generations of epimutations (Bewick *et al*. 2016), suggesting that expression was not sufficient to drive gbM reestablishment, at least not within a few generations. However, Zilberman *et al*. (2007) found that both methylated and unmethylated genes were upregulated in *met1* mutants using microarray data, suggesting *met1* methylation mutants may have unanticipated global expression effects that make them a poor system for studying the association between gbM and expression.

Another approach to test for associations between gbM and expression has been comparative genomics, which has the advantage of integrating effects over evolutionary time. Here again the results have been inconsistent. For example, Bewick *et al*. (2016) and Bewick *et al*. (2019) found no effect of the loss of gbM on gene expression in *E. salsugineum* compared to *A. thaliana*. In contrast, Muyle and Gaut (2019) found a small but significant decrease in expression associated with genes that lost gbM in *E. salsugineum*, based on a reanalysis of the data from Bewick *et al*. (2016). In another effort, Takuno *et al*. (2017) identified genes that changed methylation status between *A. thaliana* and *Arabidopsis lyrata*. They found a trend: genes that had gained gbM between species tended to also shift toward higher expression levels. Finally, Seymour and Gaut (2019) studied eight grass species and found that genes that were gbM in all eight species tended to have higher and less variable expression, although the effect is small. This last observation is consistent with previous observations that gbM is associated with less variable gene expression both within and between species (Zilberman *et al*. 2008; Coleman-Derr and Zilberman 2012; Steige *et al*. 2017; Takuno *et al*. 2017; Horvath *et al*. 2019; Seymour and Gaut 2019), suggesting it has a homeostatic effect on expression (Zilberman 2017).

In addition to a potential – but unresolved – association with gene expression, a second hypothesis of gbM function is that it prevents aberrant internal and/or antisense transcription (Tran *et al*. 2005; Maunakea *et al*. 2010). Here again the evidence is unclear, because studies comparing gbM mutants to wild type mouse embryonic stem cells have been contradictory (Neri *et al*. 2017; Teissandier and Bourc’his 2017). In plants, Bewick et al (2016) found no evidence that gbM prevents antisense transcription in *met1* derived regions of *A. thaliana* epiRILs compared to orthologous wild type regions. However, Choi et al (2020) has shown that gbM and histone H1 jointly suppress antisense transcription in a comparison of *met1,h1* double mutants to WT *A. thaliana*.

The third hypothesis is that gbM improves splicing fidelity and prevents intron retention. There is some evidence for this hypothesis, because the alteration of DNA methylation impacts alternative splicing in honey bee and mouse embryonic stem cells (Li-Byarlay *et al*. 2013; Yearim *et al*. 2015). Horvath *et al*. (2019) has found evidence to support this hypothesis by comparing gbM genes to unmethylated genes in *A. thaliana*, but Bewick *et al*. (2016) found no evidence for this effect by comparing *met1* epiRILs to wild type plants. Overall, the contradictory findings regarding the possible function of gbM suggests that its effects, if any, must be relatively small.

While assays of the functional relevance of gbM have provided mixed results, evolutionary patterns of gbM have provided consistent but indirect evidence of its potential importance. Across plant species, gbM genes are generally longer, enriched for housekeeping and other important functions and evolve more slowly than unmethylated genes (Takuno and Gaut 2012, 2013; Takuno *et al*. 2017; Seymour and Gaut 2019). Moreover, comparative analyses have shown that gbM is conserved for orthologous genes between species as distantly related as ferns and angiosperms (Takuno and Gaut 2013; Seymour *et al*. 2014; Takuno *et al*. 2016; Niederhuth *et al*. 2016; Seymour and Gaut 2019). This last characteristic of gbM is surprising because DNA methylation is mutagenic and elevates C to T substitutions (Bird 1980). Hence, the conservation of gbM over millions of years suggests that the mutagenic feature of methylation is counterbalanced by an advantageous effect that acts to maintain gbM in specific genes (Zilberman 2017). However, another possible explanation for the strong conservation of gbM within a specific set of genes is that *de novo* methylation biases, such as those that target the CHG-gain genes of *E. salsugineum* (Wendte *et al*. 2019), have been conserved across species over vast periods of evolutionary time.

Clearly several questions about gbM function and evolution remain unresolved. Here we move away from experiments and comparative studies and employ population genetic approaches to study gbM. Thus far, the tools of population genetics have been applied to epigenetic phenomena in only a handful of studies. For example, van der Graaf *et al*. (2015) found similar epimutation rates between *A. thaliana* populations and > 31 generations of epimutation accumulation lines, suggesting that selection has not impacted global patterns of CG methylation diversity in that species. They nonetheless argued, based on the rate of epimutation events, that selection of epiallelic states could be an important process. Wang and Fan (2014) developed a modification of Tajima’s D for application to methylation data and used it to demonstrate that new genes have an excess of rare epialleles, which they interpreted was consistent with directional selection on an epigenetic state. Two other studies have used site frequency spectra (SFS) to test for selection on methylation data. In the first, Vidalis *et al*. (2016) estimated the SFS of cytosine sites within genes of a sample of 92 *A. thaliana* individuals, but they did not detect a deviation from neutrality. More recently, studies have hinted at selection on methylation, because an SFS analysis at the level of 100bp regions detected weak but significant selection on methylation levels (Xu *et al*. 2020) and because germline promoter methylation was inferred to be deleterious in humans (Boukas *et al*. 2020).

Here we extend the SFS approach to data from the 1001 methylomes project in *A. thaliana* (Kawakatsu *et al*. 2016), to test two features of gbM. The first is whether there is evidence that gbM is subject to selection. To do so, we focus on the methylation state of genes, rather than individual sites. We focus on genes because previous work has shown that methylation is evolutionary conserved at the level of genes and not within individual sites, suggesting that the methylation state of a gene region could be the unit under selection (Takuno and Gaut 2013). Consistent with this hypothesis, Vidalis *et al*. (2016) found no evidence of selection at the cytosine level. The second is that we provide an intraspecific test of the association of gbM and gene expression by comparing the methylation state of alleles to their level and variability in expression. By harnessing the power of an extensive *A. thaliana* data set, we uncover new information on the evolutionary forces that may act on epigenetic phenomena and the potential functional significance of gbM.

## Materials and Methods

### Datasets

Methylation and expression files for *A. thaliana* from Kawakatsu *et al*. (2016) were downloaded from GEO (accessions GSE43857, GSE80744, GSE54292, GSE43858, GSE54680). The files consisted of tables with one line per cytosine showing the number of methylated and unmethylated bisulfite sequencing (BS-seq) reads for each methylome, and tables with one line per gene showing the number of reads mapping for each transcriptome. The dataset included 1211 samples sequenced by BS-seq and 1195 by RNA-seq. More precisely, 927 *A. thaliana* were grown at 22°C and their methylomes were sequenced by BS-seq at the SALK Institute (Kawakatsu *et al*. 2016), of which 876 came from leaves and 51 from flower buds (with only a partial overlap in accessions between the two tissues). 144 samples had their leaf transcriptome profiled with the SOLiD system (Schmitz *et al*. 2013), and 728 samples had their leaf transcriptome sequenced by Illumina RNA-seq (Kawakatsu *et al*. 2016). Swedish accessions had their leaf BS-seq data generated at the Gregor Mendel Institute (GMI) (Dubin *et al*. 2015). These included 152 accessions that were grown at 10°C and another 121 accessions grown at 16°C. Some accessions had replicates sequenced, resulting in a total of 284 methylomes from GMI. These had corresponding leaf transcriptome data from 160 accessions grown at 10°C and from 163 accessions grown at 16°C, for a total of 323 samples sequenced by Illumina RNA-seq (Dubin *et al*. 2015). However, we detected a strong Institute-of-origin effect in the data (see Results), leading us to focus most of our analyses on leaf data from the Salk Institute (876 accessions).

To have outgroup data, we retrieved *A. lyrata* MN47 and *Capsella rubella* MTE ∼10 day old seedling shoot methylation files (Seymour *et al*. 2014). For each species, two replicates grown at 23°C were used.

### Inference of cytosine methylation

Cytosine methylation calls were already in the downloaded files from the Salk institute, and these calls were based on the method of Kawakatsu *et al*. (2016). For *A. thaliana* data from GMI (273 samples plus 11 replicates) as well as for *A. lyrata* and *C. rubella* data, we inferred cytosine methylation using the same method. Briefly, methylation was inferred for each site by performing a binomial test on the number of methylated and unmethylated reads, while taking into account the non-conversion rate (Lister *et al*. 2008). For the GMI data, the average non-conversion rate of 0.0041 was used for all samples (Dubin *et al*. 2015). P-values were corrected for multiple tests using Benjamini and Hochberg correction. Sites with ≤ 2 reads were considered as unmethylated, and sites with a corrected p-value under 0.001 were considered to be methylated.

### Inference of gene body methylation

For each gene, the methylation state was inferred using data from coding sequences (CDS), which included exons but excluded both untranslated terminal regions and introns. We used the annotation of the longest transcript to define the CDS. For each accession separately, we computed an expected methylation rate for each context (CG, CHG, CHH) across all CDSs annotated in the genome, and we used binomial tests to assess whether gene CDSs had a significantly higher proportion of methylated cytosines than the genome-wide background level of CDS methylation (Takuno and Gaut 2012). This was performed for each accession and cytosine context separately. P-values were corrected for multiple tests using the Benjamini and Hochberg correction for each accession separately.

Given the binomial results, a gene within an accession was inferred to be **gene body methylated (gbM)** if it had more than 20 CG sites and if CG methylation was significantly higher than the background (one-sided binomial p-value lower than 0.05) and CHG and CHH methylation were not significantly higher than the background (one-sided p-values higher than 0.05). A gene was inferred to be **CHG methylated** if it had more than 20 CHG sites and if CHG methylation was higher than the background (one-sided p-value lower than 0.05) and CHH methylation was not significantly higher than the background (one-sided p-values higher than 0.05). CHG methylated genes also tended to be CG methylated, but CG methylation was not required in our categorization. A gene was inferred to be **CHH methylated** if it had more than 20 CHH sites and if CHH methylation was higher than the background (one-sided p-value lower than 0.05). CHH methylated genes also tend to be CG and CHG methylated. Finally, a gene was inferred to be **unmethylated (UM)** if it had more than 20 CG sites and if CG, CHG and CHH methylation were not significantly higher than the background (one-sided p-value higher than 0.05). In any other case, the gene methylation state was not inferred. Altogether, by applying this approach, we identified the frequency of methylation states across alleles among 1211 accessions and for ∼27,000 genes.

### Inference of ancestral methylation state

For each gene, the ancestral methylation state in *A. thaliana* was inferred using methylation data from *A. lyrata* and *C. rubella*. To this end, we used the CoGe tool SynMap3D (Lyons and Freeling 2008) to infer orthologous syntelogs among *A. thaliana, A. lyrata* and *C. rubella*. We differentiated between orthologs and out-paralogs (paralogs caused by duplications that predate speciation) using pairwise dS values between syntelogs. Based on the distribution of dS values (Supplementary Figure S1), log10(dS) values were filtered to be lower than -0.39 for all species pairwise comparisons, which is equivalent to dS values lower than 0.407. After this filtering, 14,718 orthologous syntelogs were identified among the three species.

Two shoot replicates grown at 23°C were available for each outgroup species (*A. lyrata* and *C. rubella*) (Seymour *et al*. 2014). For every gene, the ancestral methylation state was inferred as the shared state between the two outgroups and their replicates. If the two replicates of a species had different methylation states for a gene, or if the gene had different methylation states between *A. lyrata* and *C. rubella*, we excluded it from analyses as having an ambiguous ancestral state.

### Inference of genes undergoing CG methylation epimutations

We also investigated the set of CHG-gain genes from *E. salsugineum CMT3* overexpressing transgenic lines by retrieving the list of 8,704 CHG-gain genes from Wendte *et al*. (2019). The best blast hit – as provided in the genome reference – was used to infer the ortholog in *A. thaliana* for 8,025 of these CHG-gain genes.

### Site frequency spectrum (SFS)

The unfolded SFS was drawn for two gene methylation states, gbM and UM. mCHG and mCHH states were excluded from the SFS (see Supplementary Figure S2 for the distribution of the proportion of mCHG and mCHH accessions across all genes). Genes were included in the SFS only when methylation status could be determined in 200 or more accessions, and genes that had over 70% of accessions with mCHG or mCHH methylation state were discarded as possibly being pseudogenes or misannotated TEs.

The number of accessions with an inferred methylation state *n* varied among genes due to missing data, so that the site frequency spectrum sample size varied among genes. To cope with this missing data, we defined *n’*, the minimum required number of accessions with characterized methylation, and applied a hypergeometric projection of the observed SFS into a subsample of size *n’*=200. Genes sampled in less than *n’* accessions were discarded. The frequency of the derived allele in the reduced sample follows a hypergeometric distribution. Given *k* the frequency of the derived allele in the original sample of size *n*, the probability that *i* copies were observed in the reduced sample of size *n’* is (Hernandez *et al*. 2007):

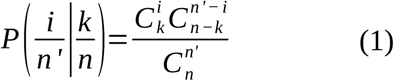

### Estimation of selection using the site frequency spectrum

Given the SFS, we estimated the strength of selection acting on methylation variants using the model of Charlesworth and Jain (2014). The model was designed to characterize the evolutionary forces acting on epigenetic markers, which evolve at much higher rates than DNA sequences when single sites are considered (Becker *et al*. 2011; Schmitz *et al*. 2011). We adapted the model for application to our biological question of whether selection acts on gene methylation states. Genes can either be gbM or UM, with *μ* and *ν* the mutation rates from one state to the other:

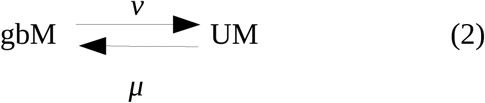

The model assumes a randomly mating diploid population of constant effective population size *N*_*e*_ which is at mutation-selection equilibrium. The model further assumes that alleles are semi-dominant and that sites are independent. We estimated *N*_*e*_ using available polymorphism measures in *A. thaliana* (Alonso-Blanco *et al*. 2016): 10,707,430 total SNPs were detected in 1135 genomes of size 135Mb, resulting in a Watterson theta *θ*_*w*_=0.00955 (Charlesworth and Charlesworth 2010). Using a mutation rate μ=7.10^−9^ (Ossowski *et al*. 2010) and *θ*_*w*_=4*N*_*e*_*μ*, we estimated *N*_*e*_*≈*341,000. These values were similar to previous diversity measurements in *A. thaliana*, where intronic *θ*_*w*_ was estimated to be 0.0082 (Nordborg *et al*. 2005), but the actual *θ*_*w*_ may be higher due to biases in its estimation (Korunes and Samuk 2020).

If the UM state is advantageous over the gbM state, the probability that a sample of *n* individuals segregates for *k* UM variants and *(n-k)* gbM variants at a given gene is (Charlesworth and Jain 2014):

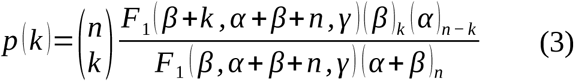

Where *F*_*1*_ is the confluent hypergeometric function, *(x)*_*n*_ is Pochhammer’s symbol, *α=4NN*_*e*_*μ, β=4NN*_*e*_*ν* and *γ=4NN*_*e*_*s*_*UM*_ with *s*_*UM*_ the selective advantage of the UM methylation state over gbM. The model can easily be adapted to a case where the gbM state is advantageous over the UM state by switching *α* and *β* in equation (3) and defining *s*_*gbM*_ the selective advantage of the gbM state.

The likelihood of the model is:

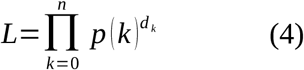

Where *d*_*k*_ is he number of genes observed with *k* UM accessions and *(n-k)* gbM accessions. Parameters of the model *μ, ν* and *s*_*UM*_ (or *s*_*gbM*_) were estimated using a Markov Chain Monte Carlo (MCMC) random walk with 100,000 generations as in Xu *et al*. (2020). The first 25% of MCMC generations were removed as burn-in. Parameters were sampled every 100 generations, providing around 750 samples for the posterior distributions of parameters. The lambda parameters for scale proposal distribution were adjusted to obtain parameter acceptance rates between 20% and 70%. Both segregating and fixed sites of the SFS were used in the model. Final parameter values were obtained from the mean of the posterior distribution and the credible interval from the 95% margins of the posterior distribution. We ran the algorithm three times with random starting points to ensure that the global maximum was found. In order to test whether selection acting on the UM or the gbM state was significant, we compared the previous full model to a reduced model where the selection coefficient was fixed to zero using a likelihood ratio test with degree of freedom 1. For each run, the expected SFS (using inferred parameter values from the best model) was compared to the observed SFS using a Pearson’s *χ*-square test in R.

### Statistical study of the link between gbM and gene expression level

We measured the effect of gbM on expression level using the Salk Institute leaf dataset, which consisted of 679 accessions with both leaf methylation data and leaf expression data in the form of raw RNA-seq read counts. This number of accessions differed from the previous 876 Salk accessions used for the SFS analysis due to missing leaf expression data for some accessions. We constructed a linear model with mixed effects (equation 5) to examine the data, which was run with the R package lme4 (Bates *et al*. 2015). We did not normalize gene expression and used raw read numbers, but results were equivalent when using normalized read numbers as provided in GEO expression files. The aim of the model was to test, within each gene, for an association between a change in gene methylation state and gene expression across *A. thaliana* samples. To account for expression variability among genes, the model incorporated a random gene effect (see equation 5). The random gene effect captures variability in gene expression due to average differences among genes. We also defined a fixed effect called gene methylation state (equation 5), which consists of the states described above (e.g., gbM, mCHG, mCHH and UM) and applies to each gene epiallelic state within each accession. Significance for the fixed effect was determined by comparing the fit of the full model to a nested model without the fixed effect, using the anova function in R. Expression level was measured as raw read counts and log transformed. The R package lsmeans (Lenth 2016) was used to estimate pairwise differences between each pair of methylation states (i.e. gbM versus UM, UM versus mCHG etc.).

Our linear model can be expressed as:

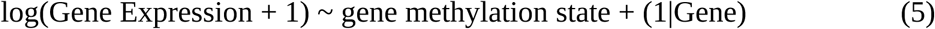

We also developed two linear mixed-effects models to investigate the potential relationship between genetic and epigenetic states of alleles. The model included the number of CG dinucleotides (#CG) and the epiallelic methylation state, as fixed effects, and the random gene effect:

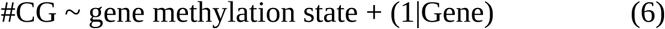

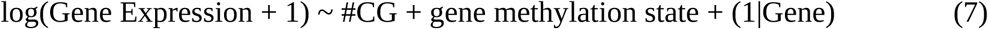

## Results

### Detection of selection acting on gene methylation level

We used publicly available methylation datasets (Seymour *et al*. 2014; Kawakatsu *et al*. 2016) to infer the methylation state of genes in *A. thaliana* accessions and two closely related outgroups. We first recognized that BS-seq sequencing of the *A. thaliana* 1001 methylomes was carried out by two research Institutes (Salk and GMI), and so we compared methylation patterns and levels between the two institutes. We found that the global rate of CHH methylation was significantly higher in accessions sequenced by GMI (2.3%) compared to the Salk Institute (0.28%, Supplementary Figure S3), regardless of the geographic origin of accessions (Supplementary Figure S4). This heterogeneity in the raw data had the potential to impact downstream gene methylation inferences (Supplementary Figure S5). We therefore focused on a single source – i.e, the Salk data – because it had the highest number of samples (927 methylomes). For similar reasons, we also focused only on methylome data from a single tissue (leaf), leading to total analysis sample of 876 accessions.

Given the data, we inferred the gbM status for each gene in each individual separately to calculate the unfolded SFS for two gene methylation states - gbM and UM – after downsampling to 200 accessions (see Material and Methods for details). Altogether, we plotted the SFS based on 23,868 genes and found that, after downsampling to 200 accessions, most genes were fixed for the UM methylation state in *A. thaliana* wild populations (11,901 genes), but there was also a subset of 983 genes fixed for gbM alleles (Figure 1.A). Given the inferred SFS, we applied the model of Charlesworth and Jain (2014) to infer the selection coefficient, *s* in a model where the UM state is advantageous and in another model for which the gbM state is advantageous. Based on all 23,868 genes together, we found that the model that best fit the data was one where the UM state is advantageous (*s*_*UM*_=8.68.10^−8^), with a *p*-value of 0.0086 based on comparing the two models with selection to a model without selection (Table 1). Note that the expected SFS based on estimated parameters fit the observed SFS quite well (Figure 1.A), with no significant difference between them (Pearson’s *χ*-squared test p=0.343).

**Table 1:**
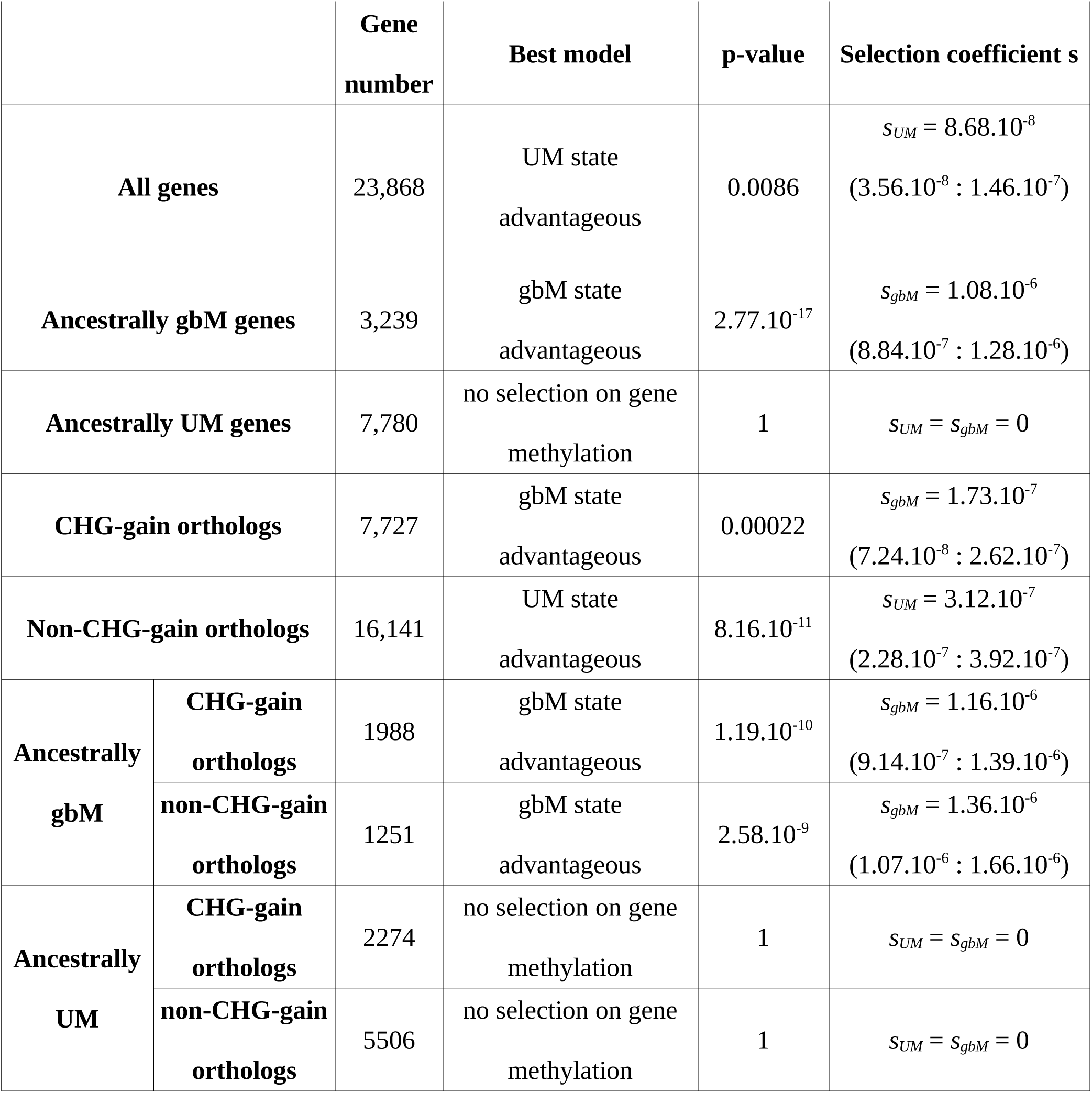
Estimation of selection acting on gene methylation state. A likelihood-based approach was used to infer the best model to explain the SFS (see Materials and Methods for details). The p-value shows the result of the likelihood ratio test between the model with selection and the neutral model (s=0). The estimated selection coefficient *s* is shown with a 95% credible interval in parenthesis. Details on inferred values of other parameters of the model can be found in Supplementary Table S1. These results come from one MCMC run and are equivalent to results obtained from two independent runs with random parameter initiation values (Supplementary Table S1).

**Figure 1:**
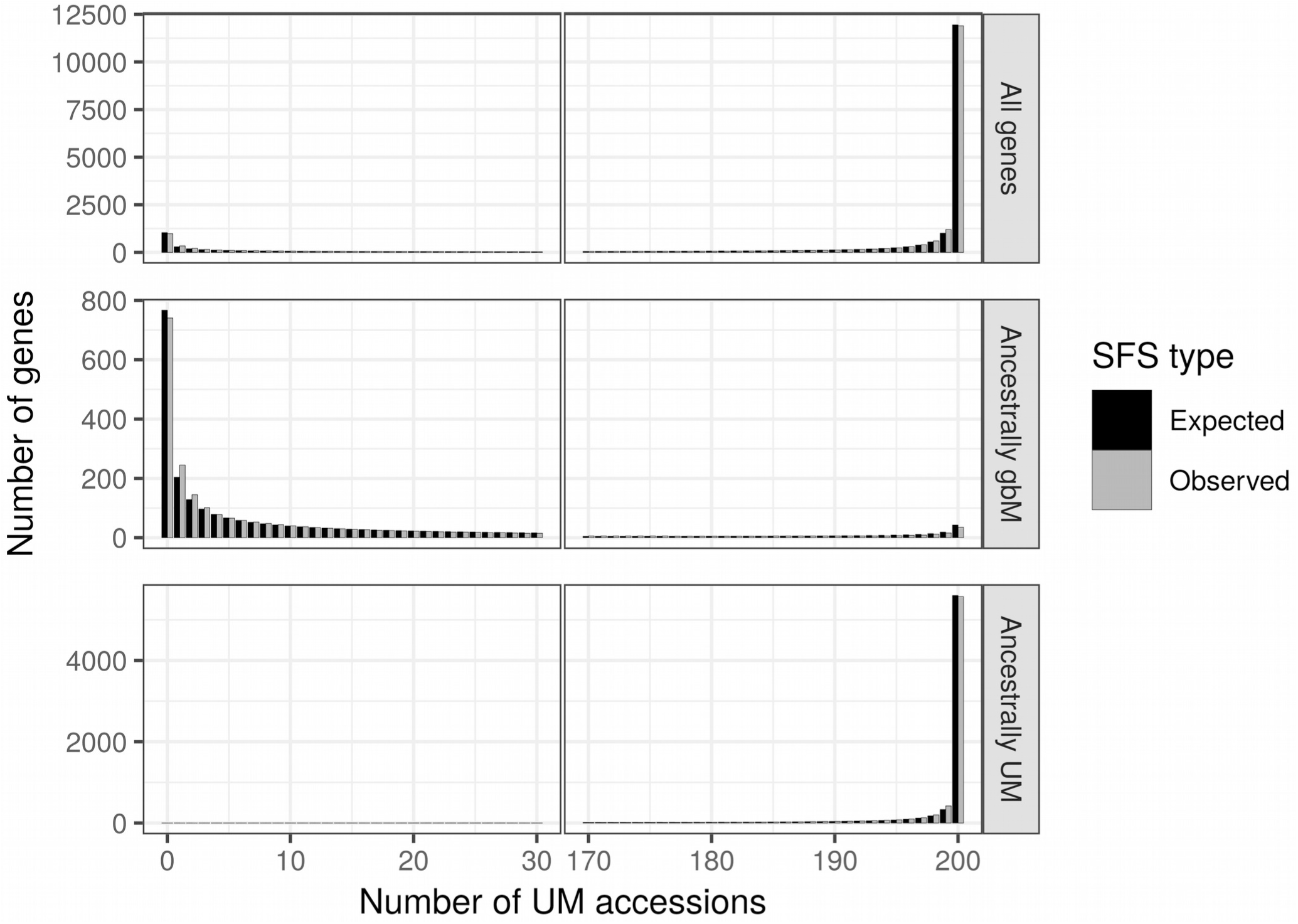
Expected and Observed Site Frequency Spectra (SFS) of gene body methylation. The x-axis provides the number of UM accessions, out of a sample of 200. For these data, accessions that are not UM are gbM, meaning that genes with 200 UM accessions are fixed for the UM state in *A. thaliana* and genes with 0 UM accessions are fixed for the gbM state. The number of genes is provided on the y-axis. **A**. All genes (23,868 genes), **B**. ancestrally gbM genes (3,239 genes) and **C**. ancestrally UM genes (7,780 genes). For visualization purposes, a gap was introduced in the x-axis. The expected SFS were drawn using the parameters estimated by the mcmc, using the best model in Table 1. All three expected SFS fit the observed SFS well and did not differ significantly from the observed distribution (Pearson’s *χ*-squared test p>0.4).

Our SFS illustrates that gbM is markedly bi-modal, which is consistent with the fact that gbM is associated with a finite but conserved set of orthologous genes across angiosperms (Takuno *et al*. 2016). It seems reasonable to presume, then, that genes that are evolutionary conserved as gbM may be under different selection regimes than those that are evolutionary conserved as UM. Accordingly, we repeated analyses after splitting ancestrally gbM and UM genes (Figures 1.B and 1.C). To infer the ancestral state, we used two outgroups (*A. lyrata* and *C. rubella*), identified syntelogs for 14,718 genes among the three species, and then inferred the ancestral methylation state by parsimony when both outgroups and their replicates had the same methylation state. After excluding 3,699 genes for either missing methylation state inferences or for having ambiguous ancestral state, we applied the model to a set of 3,239 genes that were inferred to be ancestrally gbM, estimating a small (*s*_*gbM*_ = 1.08.10^−6^) but highly significant (*p* = 2.77.10^−17^) selection coefficient (Table 1). This result implies that there is weak but detectable selection to maintain methylated alleles within genes that are ancestrally gbM. In contrast, 7,780 ancestrally UM genes were estimated to be under no selection to retain UM nor gbM alleles in *A. thaliana* wild populations (*s*_*UM*_ = *s*_*gbM*_ = 0, Table 1).

We have mentioned both that there is no gbM in *E. salsugineum* due to the loss of *CMT3* (Bewick *et al*. 2016) and that complementation of *E. salsugineum* with a functional copy of *A. thaliana CMT3* leads to the accumulation of DNA methylation in ‘CHG-gain’ genes (Wendte *et al*. 2019). These results suggest that DNA methylation epimutation rates are not homogeneous among genes, which could be problematic for our model. We therefore repeated the previous analyses separately for CHG-gain and for non-CHG-gain genes in *A. thaliana*, based on identifying the orthologs of CHG-gain genes from *E. salsugineum* (Materials and Methods). We found that 7,727 CHG-gain genes were under significant selection for retaining the gbM state (Table 1), again implying that selection acts on gene methylation state. On the other hand, the set of 16,141 no-CHG-gain genes were estimated to be under selection to retain UM status. The model also estimates epimutation rates; consistent with implications of the *E. salsugineum* experiment (Wendte *et al*. 2019), we found that the mutation rate *m* from the UM state to the gbM state was ∼1.77 times higher in CHG-gain compared to no-CHG-gain genes. In contrast, the epimutation rate *v* from gbM to UM was ∼0.79 lower in CHG-gain compared to no-CHG-gain genes (Supplementary Table 1).

Finally, we repeated the analyses after splitting CHG-gain and non-CHG-gain genes into ancestrally gbM and ancestrally UM genes, because we have shown that the SFS of both features (methylation state and CHG-gain state) suggest ongoing selection. Ancestrally gbM genes were under significant selection to remain gbM, regardless of whether they were targeted by additional methylation epimutations (CHG-gain) or not (non-CHG-gain, Table 1). However, ancestrally UM genes were under no significant selection to remain UM or become gbM (Table 1). Our results were therefore confirmed after splitting the dataset into sets of genes with putatively homogeneous epimutation rates; overall they provide evidence that the methylation state of alleles is associated with natural selection.

### Effect of gene methylation state on gene expression level in *A. thaliana* wild populations

*E. salsugineum* lacks gbM due to the loss of CMT3 (Bewick *et al*. 2016), but there has been some debate about the effects of this gbM loss on gene expression. Bewick et al (2016) found no effect, a result upheld by later analyses (Bewick *et al*. 2019). However, using different statistical approaches, Muyle and Gaut (2019) found evidence for a small but significant decrease in expression level for *E. salsugin*eum genes that had lost gbM relative to the same gbM genes in *A. thaliana*. We further investigated the possible association between gbM and gene expression by analyzing expression levels from the *A. thaliana* 1001 methylome data. To make this assessment, we focused on leaf expression and methylation data from the Salk Institute for 679 accessions and 23,261 genes with polymorphic methylation states (i.e., genes fixed for a given methylation state were removed from consideration). The availability of these data permitted a test of whether epiallelic methylation states are associated with differences in expression.

We analyzed the data using a linear model with mixed effects (equation 5, Materials and Methods). The model was written to measure within gene expression variation, and then test for a significant effect of methylation state across all genes. This approach is possible due to polymorphisms in gene epiallelic states (or epialleles) among accessions. Treating genes as random effects and epiallele state (gbM, UM, mCHG or mCHH) as a fixed effect, we found that epiallele methylation state had a significant effect on gene expression level (*χ*^2^ = 19,300 and p-value = 0 when comparing a linear model with and without gene methylation state effect). We repeated these analyses between pairs of methylation states, comparing along an expected hierarchy of expression levels defined as gbM > UM > mCHG > mCHH. Our results confirmed that expected hierarchy, because gene expression in accessions that had the gbM epiallelic state was significantly higher than for accessions that had the UM epiallelic state for that same gene, globally across all genes (Figure 2). Similarly, we found that UM alleles had higher expression than mCHG alleles and that mCHG alleles were more highly expressed that mCHH alleles (Table 2). Altogether, these results show that within a gene, an accession with the gbM epiallelic state is consistently associated with the highest gene expression level, while the mCHH state is consistently associated with the lowest gene expression level. However, the estimated differences in expression levels were very small (0.0563 log read count difference on average between gbM and UM methylation states, Table 2, which is equivalent to 1.058 raw read count difference on average). It is important to note that linear models can detect small mean differences as significant so long as those differences are prevalent across the entire dataset.

**Table 2:** Pairwise comparison of the effect of gene methylation state on gene expression level in *A. thaliana* 1001 methylome data. A generalized linear model with mixed effects was used to estimate the effect of gene methylation state on gene expression (see Materials and Methods for details). Gene expression was measured as raw read counts and log transformed, but the results were equivalent when performed on normalized read counts. The table shows the average differences in log expression levels between pairs of methylation states (estimates) and their associated standard error, t-ratio and p-value after correction for multiple tests. For example, the gbM state is consistently associated with a 0.0563 higher log read count compared to the UM state, on average across all genes and accessions.

| Contrast | Estimate | Standard Error | z-ratio | p-value |
| --- | --- | --- | --- | --- |
| gbM – UM | 0.0563 | 0.0011 | 51.56 | 0 |
| UM – mCHG | 0.1093 | 0.0033 | 33.13 | 0 |
| mCHG – mCHH | 0.2275 | 0.00359 | 63.33 | 0 |

**Figure 2:**
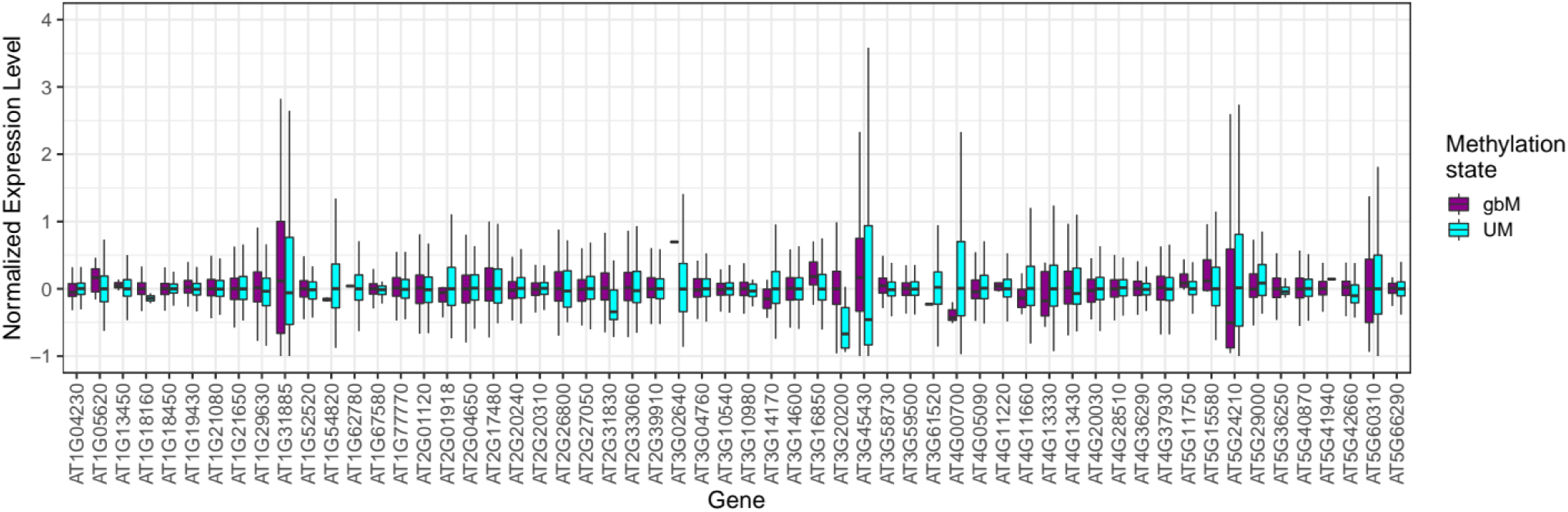
Normalized expression levels in a random set of genes with both UM and gbM epialleles. To make this figure, a random set of 100 genes was selected and only genes with both UM and gbM epialleles were retained. Genes with median read number under 10 were also excluded, leaving 57 genes. For each gene, the box plots indicate normalized expression levels for gbM and UM epialleles, with the lines representing the medians and the width of the boxplot the 1^st^ and 3^rd^ quartile. The figure shows that accessions that have a gbM epiallele tend to have a higher median normalized expression compared to accessions that have a UM epiallele, because 34 out of 57 genes in the represented random gene set have higher median expression levels for gbM epialleles. Although the differences are small and some genes show the opposite pattern, the overall effect is significant in a linear model across genes (see text for details). For each accession, normalized expression level was computed as follow: (accession normalized read number – median gene normalized read number) / median gene normalized read number.

In order to further test the robustness of our results, we performed two additional analyses. First, to confirm that the results were not an artifact of the linear model, we reran the model after randomly permuting methylation states without replacement among accessions and genes. These permutations removed associations between methylation states of an allele and their expression, and hence we did not expect to detect significant effects with permuted data. We ran the model on 1001 permuted datasets. As expected, the correlation between gene methylation state and expression was significant at *α* = 0.05 only ∼5.0% of the time, because we detected significance in 54 of 1001 permutations (Figure 3). What is more, the *p*-value obtained from the real dataset was more than 8 orders of magnitude lower than the lowest *p*-value obtained on any of the permuted datasets. These permutation results illustrate both that the model is well-behaved and that the observed data are strongly unexpected under a null hypothesis in which methylation and expression are not linked (Figure 3). Second, we compared expression levels between UM and gbM alleles within single genes for the 11,613 genes that had at least one UM accession and one gbM accession. We found accessions with gbM epialleles had a higher median expression level than accessions with the UM state for 6,122 out of 11,613 (or 52.72% of genes). This proportion represents a significant deviation (binomial test, two-sided, *p*-value 5.10^−9^) from the the expected value of 50% under the null hypothesis that epiallelic state does not affect expression.

**Figure 3:**
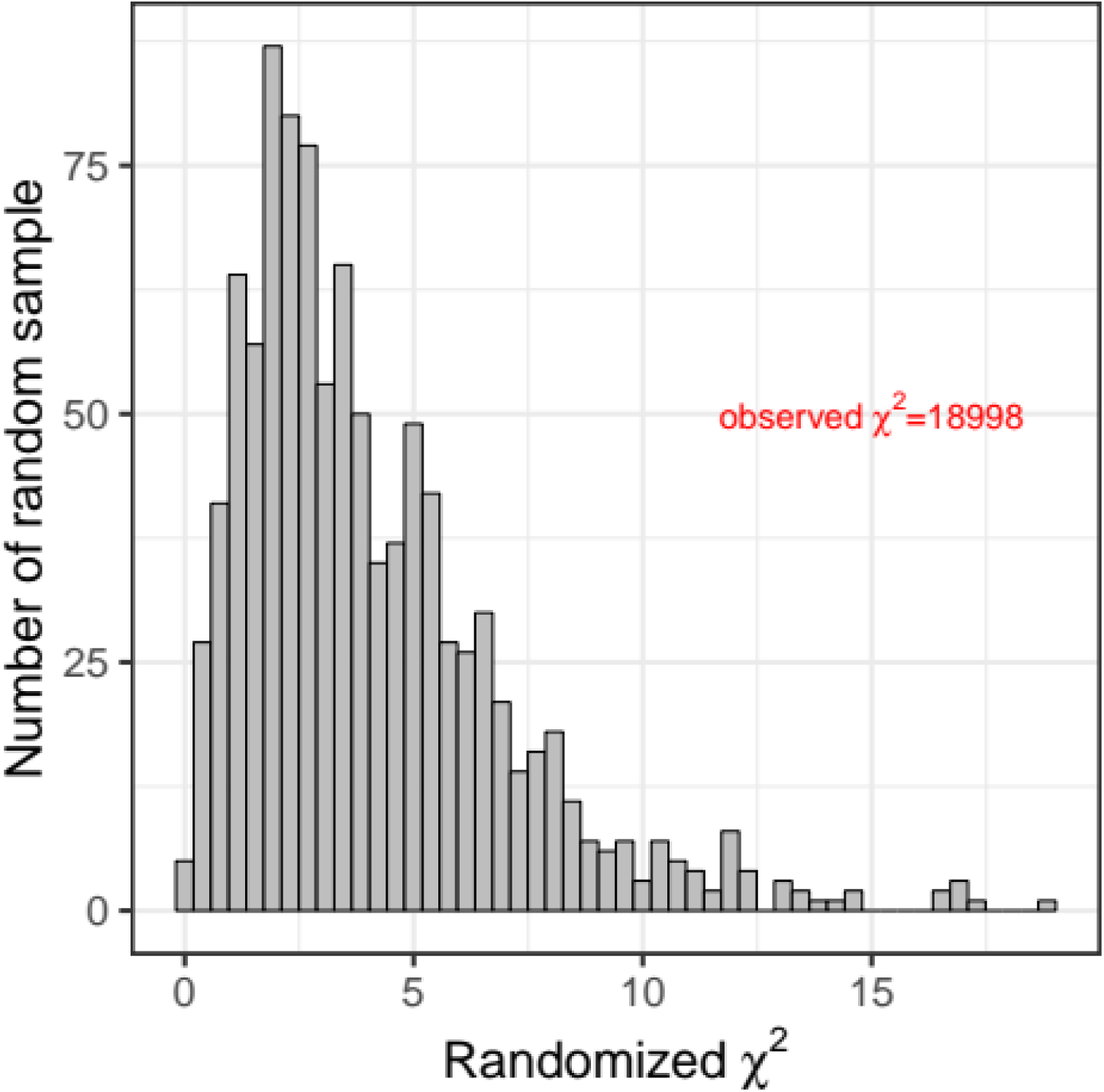
Distribution of *χ*^2^ values obtained after randomizing the data to test the association between gene methylation and expression level in the *A. thaliana*. For each permuted data set, a linear model with mixed effects was used to assess the correlation between methylation and expression levels (Materials and Methods, equation 5). A *χ*^2^ value > 7.8 represents a significant correlation between methylation level and expression level within genes (i.e., a *p*-value < 0.05 with 3 degrees of freedom), and such values were observed 5.4% of the time. The observed *χ*^2^ (18,998) was higher than the highest *χ*^2^ obtained on randomized data (18.94) by over a thousand-fold.

These data also present the opportunity to test the homeostasis hypothesis, which posits that gbM acts to stabilize gene expression (Zilberman 2017). Under this hypothesis, we expect the coefficient of variation to be lower across gbM epialleles than for UM epialleles. To test the hypothesis, we focused on the 10,327 genes that harbored at least two accessions of both UM and gbM epiallelic states in the population and then controlled for sample size differences by randomly sampling the same number of accessions for both epialleles within each gene. We found that the gbM state had a significantly lower coefficient of variation than the UM state (one-sided Wilcoxon paired signed rank test p = 4.77.10^−6^). Alternatively, we simply counted the number of genes that had a higher coefficient of variation for the UM state compared to the gbM state; 5,357 (or 61.56%) genes had more variable expression among UM epialleles, representing a highly significant deviation from the null expectation of 50% (binomial test, two-sided, *p*-value 1.5.10^−4^).

## Discussion

We have utilized the dataset of 1001 *A. thaliana* methylomes (Kawakatsu *et al*. 2016) to examine features of the population genomics of epiallelic states, with a focus on gbM. Our analyses reveal two main observations. The first is that the SFS of allelic states yields information about selection. We find that all genes taken together are under selection to be UM, however, ancestrally gbM in *A. thaliana* are under selection to remain gbM. The second observation is that analysis of this extensive dataset has revealed an association between epiallelic state and patterns of expression, in terms of both expression level and stability. Both observations have broad relevance but also have caveats that must be considered.

### gbM is under selection in *A. thaliana*

This study relies on 876 leaf methylomes in *A. thaliana* to investigate whether gbM is under selection and also on an SFS approach to detect selection that is specifically designed for epigenomic data (Charlesworth and Jain 2014). Our work offers the first evidence that ancestrally gbM genes are subject to natural selection to remain gbM. As selection only acts on traits that impact fitness, these results suggest that gbM has a function, at least in *A. thaliana* and maybe in other organisms that have similar gene methylation patterns (i.e., plants, mammals and insects).

The estimated selection coefficient for the advantage of gbM, *s*_*gbM*_, is small (1.10^−6^ on average, Table 1), resulting in *γ=4NN*_*e*_*s*_*gbM*_=1.38. Values of *4NN*_*e*_*s* lower than 1.0 are typically considered neutral (Charlesworth and Charlesworth 2010). The inferred values of selection coefficients acting on gbM are similar to values estimated for codon usage bias, a phenomenon known to be under weak but significant selection in species with large enough *N*_*e*_ (Galtier *et al*. 2018). For example, leucine, valine, isoleucine and arginine have estimated *γ* values for selection on codon usage between 1 and 2 in *A. lyrata* (Qiu *et al*. 2011). GbM is therefore a trait that seems to have a weak impact on fitness, but natural selection may be substantial enough to maintain it in a subset of important genes through time, just like for codon bias.

Interestingly, our results on all genes taken together were opposite to those based on gbM genes, because all genes are inferred to be under selection to be unmethylated (Table 1). These results suggest, first, that the selection on gbM state varies among genes and, second, that gbM might be associated with a selective trade-off (Kiefer *et al*. 2019). That is, gbM is advantageous for some genes, but can also be deleterious, perhaps due to the increased mutation rate on methylated cytosines, energetic costs or effects on chromatin structure. We therefore argue that the advantages of gbM outweigh its putatively deleterious mutagenic effects in a subset of genes, but in other genes either gbM offers no advantage or the advantage is not strong enough to compensate for higher mutation rates.

Our inferences are of course subject to caveats. The first set of caveats is related to our application of the model of Charlesworth and Jain (2014), which assumes uniform epimutation rates across the entire genome. To help address this limitation of the model, we investigated ancestrally UM and gbM genes separately, and we also separated the set of CHG-gain genes identified in *E. salsugineum* from the non-CHG-gain genes. We separated the latter two because CHG-gain genes may have elevated rates of *de novo* epimutation (Wendte *et al*. 2019); indeed, we estimate that CHG-gain genes have 1.77-fold higher mutation rates from the UM to gbM state. However, the datasets all inferred similar trends, including estimates of *s* that were significantly different from zero, with selection to retain methylation for ancestrally gbM genes (Table 1). These results suggest that heterogeneity in epimutation rates across genes are unlikely to drive our results, although we advocate for future investigations into CMT3 targeted genes in *A. thaliana*, because their dynamics could differ from *E. salsugineum* given the ∼47 million year (my) divergence between species (Arias *et al*. 2014). Additional limitations of the model include assumptions about outcrossing, semi-dominance, independence among sites, demographic equilibrium and mutation-selection balance (Charlesworth and Jain 2014). Clearly, the first two of these assumptions are violated by our study organism (*A. thaliana*), which is predominantly selfing. However, we treated each individual as haploid (in the sense that we did not separate the two alleles of a gene), which reduces to sampling one allele per individual from an outcrossed population. Nonetheless, we find that the model fits the data well despite these limitations (Figure 1).

Another set of limitations surround the data and our treatment of the data. For example, we used data from 10-day shoots – which are a mix of leaves, stems and leaf buds – to infer ancestral states of *A. thaliana* leaves. We therefore assume that shoots adequately reflect methylation states in leaves, an assumption that appears to be reasonable for genic methylation states across tissues of *Brachypodium distachyon* (Roessler *et al*. 2016). We also treated complete CDS regions as an epiallelic state – e.g., gbM, UM – and inferred the SFS of those states. We chose to employ this approach over other options, such as investigations of the per-cytosine SFS or the SFS of differentially methylated regions (DMRs) based on the study of Takuno and Gaut (2013). This study found that an ortholog that is gbM in one species is highly likely to be gbM in another species, even when the two species in question (in this case, rice and *Brachypodium distachyon*) diverged ∼50 my ago. The remarkable feature of this observation is that the methylation of orthologs was conserved but the methylation of individual nucleotides was not. In other words, this and subsequent studies have suggested that gbM is a property of genes, not nucleotide sites nor DMRs, which are an amorphous and often statistically problematic concept (Roessler *et al*. 2016). Given their observations, Takuno and Gaut (2013) hypothesized that gbM is a threshold character, such that the functional effects of gbM require some threshold of methylation that relies on the number and distribution of cytosines across genes. We caution that we have not explicitly tested that model here, nor inferred the properties of any threshold, but our results based on contrasting gbM and UM alleles are consistent with such a model. Our focus on genes (as opposed to nucleotide sites or DMRs) is also justified from observations about CHG-gain genes in *E. salsuginuem* (Wendte *et al*. 2019).

Finally, we focus on the use of *A. thaliana* as a study organism for studies of methylation. *A. thaliana* has been used as a model system for good reason; without its genetic tools, the pathways and mechanisms of cytosine methylation in plants would not be nearly as well understood (Law and Jacobsen 2010). Similarly, the fact that it is selfing with a small genome size makes it ideal for some applications such as population genomics and epigenomics (Alonso-Blanco *et al*. 2016; Kawakatsu *et al*. 2016), leading to the generation of unique datasets like the one we have analyzed here. However, *A. thaliana* may not be the ideal model to study methylation mutants precisely because those mutants have less phenotypic effect in *A. thaliana* than in some other plants – for example, methylation mutants are in maize often lethal (Li *et al*. 2014). Consistent with this conjecture, a previous study comparing *A. thaliana* and *A. lyrata* gene methylation states has inferred that *A. thaliana* has lost gbM three times faster than gaining it (Takuno *et al*. 2017). Our estimated values of epimutation rates on all genes (Supplementary Table S1) from gbM to UM (*ν* = 2.07.10^−7^) and from UM to gbM (*μ* = 6.17.10^−8^) exactly reiterate this three-fold difference. Thus, the growing consensus is that *A. thaliana* is losing gbM through time. We hypothesize that one reason for this is the recent shift of *A. thaliana* to an inbreeding mating system, which has reduced its effective population size (Mattila *et al*. 2020) and likely led to weaker selection on epigenetic states. The overarching – and more important – point is that *A. thaliana* is likely to be a poor model to study the evolutionary forces that act on gbM, and yet our study nonetheless detects a significant selective effect.

### gbM is associated with gene expression

As we noted in the Introduction, the question of gbM function has been raised in many studies, and gene expression has been used as the proxy for function in most of these studies. The field has thus focused on a relatively simple question: Is gbM associated with gene expression? Unfortunately, the outcome of these studies has been inconsistent, owing to a wide variety of reasons that may include that: i) the effect of gbM on expression is minor; ii) some studies are underpowered to detect such an effect, particularly over short temporal scales, iii) researchers disagree on statistical approaches, particularly whether UM genes can be utilized as a control comparison to gbM genes (Muyle and Gaut 2019; Bewick *et al*. 2019); and iv) independent epigenetic marks could have redundant functions that hide the effects of gbM loss in methylation mutants (Choi *et al*. 2020).

Our work here has, however, taken a unique approach, which is to examine the association of intraspecific variation in epialleles and expression levels across genes. This approach makes it possible to test (both within and across genes) whether a change in methylation state within the population associates with differences in expression level. To our knowledge, this is the first study to integrate intraspecific variation in methylation state with expression level in wild type plants. Our linear model consistently identified an effect of methylation state on expression, whether we investigated all of the defined states or compared pairs of states (e.g., gbM vs. UM; Table 2). The power of this approach undoubtedly comes from the extensive data generated by the 1001 methylome consortium, because the size of the estimated effect is small. In real terms, the difference between a gbM allele and a UM allele is about 1 raw sequence read, averaged over the entire data set. Nonetheless, it is clear that this result is not an artifact of the approach, because we permuted the data and found that the observed results are far more extreme (by 1000-fold) than the permuted data. In short, the evidence for the effect is strong, even though it is small. This adds to a growing number of experimental and comparative genomic approaches that point consistently to some association between gbM and expression (Zilberman *et al*. 2007, 2008; Coleman-Derr and Zilberman 2012; Steige *et al*. 2017; Takuno *et al*. 2017; Horvath *et al*. 2019; Seymour and Gaut 2019). We also show that the variation in gene expression among accessions is lower for the gbM compared to the UM epiallelic state. This is in agreement with other studies that suggest that gbM stabilizes gene expression (Zilberman *et al*. 2008; Coleman-Derr and Zilberman 2012; Steige *et al*. 2017; Takuno *et al*. 2017; Horvath *et al*. 2019; Seymour and Gaut 2019).

Our results point to selection on gbM perhaps, in part, due to its association with gene expression. But there remain two difficult questions. The first is whether selection is on gbM itself – i.e., the epigenetic states directly – or on associated factors, such as chromatin factors or even underlying sequence features that may contribute to gbM in some unknown way. Unfortunately, we find no convincing method to discriminate among an associated *versus* a direct effect of gbM, and we must thus be careful to conclude that selection acts directly on the epigenetic state. However, to investigate this question, we ran a linear model with mixed effects to study the association between the number of CG dinucleotides (#CG) and gene methylation states in the Salk Institute data (see Materials and Methods for details). There was a significant correlation between #CG and methylation states (*χ*^2^ = 17,262 and *p*-value = 0 when comparing a linear model with and without gene methylation state effect). This model also demonstrates that gbM epialleles have more CG dinucleotides than UM epialleles (linear model pairwise contrast estimate = 1.338, p < 0.0001). Surprisingly, however, when including both methylation state and #CG in a linear model to explain expression variation (see Materials and Methods), the methylation state remains the main influence on gene expression. Moreover, accessions with higher #CG are significantly less expressed (linear model estimate -5.77.10^−3^, p<2e-16), which opposes the effect of gbM on expression. Together, these analyses illustrate that the epiallelic state is not independent of the underlying sequence, as measured by #CG, but it also hints that epigenetic state contributes to phenotype in a way that is not easily explained by variation in the number of CG dinucleotides alone.

The second difficult question is function: what does gbM actually do? We cannot yet answer this question, especially given the inconsistent evidence from a variety of organisms and experiments (Zilberman *et al*. 2007, 2008; Coleman-Derr and Zilberman 2012; Li-Byarlay *et al*. 2013; Yearim *et al*. 2015; Bewick *et al*. 2016, 2019; Neri *et al*. 2017; Steige *et al*. 2017; Teissandier and Bourc’his 2017; Takuno *et al*. 2017; Horvath *et al*. 2019; Muyle and Gaut 2019; Seymour and Gaut 2019; Choi *et al*. 2020). We note, however, that histone H1 was recently shown to have a similar effect to DNA methylation in TEs and genes (Choi *et al*. 2020). In that study, expression of antisense transcripts was activated in 710 genes following methylation loss in *h1,met1* double mutants, at a level that was not positively correlated to sense transcription changes. This finding definitely demonstrates that, at least for some genes, gbM can repress antisense transcription in *A. thaliana* jointly with H1. We hypothesize that the inhibition of antisense transcription requires a threshold of cytosine methylation, which we have captured by studying the methylation state for the entire CDS. Even if this is true, there are still unanswered mechanistic questions about how the effect of gbM on anti-sense transcription affects the level and stability of expression.

## Authors contributions

AM and BSG conceived the project. AM ran the analyses. JRI provided the MCMC R code and critical ideas. DKS shared data. AM and BSG wrote the manuscript with input from all authors.

## Acknowledgements

AM is supported by an EMBO Postdoctoral Fellowship ALTF 775-2017 and by HFSPO Fellowship LT000496/2018-L. BSG is supported by NSF grant IOS-1542703. JRI is supported by NSF grant 1546719 and USDA Hatch project CA-D-PLS-2066-H. We would like to thank Mike May for help developing the MCMC approach used here.

## Data Availability

All data used in this manuscript were previously published (GEO accessions GSE43857, GSE80744, GSE54292, GSE43858, GSE54680).

